# Comparison of force fields to study the zinc-finger containing protein NPL4, a target for Antabuse in cancer therapy

**DOI:** 10.1101/2023.01.20.524865

**Authors:** Simone Scrima, Matteo Tiberti, Ulf Ryde, Matteo Lambrughi, Elena Papaleo

**Affiliations:** Cancer Structural Biology, Danish Cancer Society Research Center, 2100, Copenhagen, Denmark; Division of Theoretical Chemistry, Lund University, Chemical Centre, P. O. Box 124, SE-221 00 Lund, Sweden; Cancer Systems Biology, Section for Bioinformatics, Department of Health and Technology, Technical University of Denmark, 2800, Lyngby, Denmark

**Keywords:** Metal Ions, Molecular Dynamics, Quantum Mechanics, Copper, Metal coordination

## Abstract

All-atom molecular dynamics (MD) simulations are a powerful approach to study the structure and dynamics of proteins related to health and disease. Advances in the MD field allow modeling proteins with high accuracy. However, modeling metal ions and their interactions with proteins is still challenging for MD simulations. Over one-third of known protein structures bind metal ions and have various cellular functions, such as structural stability, catalysis, and regulation. NPL4 is a zinc-binding protein and works as a cofactor for p97, and together they regulate protein homeostasis. NPL4 is also of biomedical importance and has been proposed as the target of Antabuse, a drug recently repurposed for cancer treatment. Recent experimental studies have proposed that the Antabuse metabolites, bis- (diethyldithiocarbamate)-copper (CuET) and cupric ions released from CuET, induce NPL4 misfolding and consequent aggregation. However, the molecular details of the mechanisms of interactions of Antabuse metabolites with NPL4 and the consequent structural effects are still elusive. In this context, biomolecular simulations can help to shed light on the related structural details. To apply MD simulations to NPL4 and its interaction with copper or Antabuse metabolites the first important step is identifying a suitable force field to describe the protein in its zinc-bound states. We first examined different sets of non-bonded parameters, because we want to study the misfolding mechanism and cannot rule out that the zinc ion may detach from the protein structure during the process and copper replaces it in the metal binding site. We investigated the force-field ability to model the coordination geometry of the metal ions by comparing the results from MD simulations with optimized geometries from quantum mechanics (QM) calculations using model systems of the zinc coordination site for NPL4. Furthermore, we investigated the performance of a MD force field including bonded parameters to treat copper ions and metal-coordinating atoms in NPL4 that we obtained based on QM calculations.

## 1. Introduction

Proteins are highly dynamic entities, interconverting between different conformations, which are tightly connected to protein function and can occur on different timescales [1–5]. Structural dynamics and the related conformational changes can conveniently be modeled by molecular dynamics (MD) simulations [6–8]. Nowadays, the advances in computing architecture and algorithms for MD allow for reaching long timescales, where functionally essential protein motions may occur [9–14]. In parallel, the physical models used to describe the system in the simulation (i.e., the force fields) have witnessed a marked improvement in accuracy [15–18]. This, coupled with the development of enhanced sampling methods, allows for more statistically robust and meaningful results [19–22]. This progress has turned MD-based methodologies into a valuable toolkit to explore the mechanistic details of protein alterations due to mutations [23], post-translational modifications [24,25], or interactions with other biomolecules or metal ions [26,27].

However, the parameterization of force fields to study metal-binding proteins still represents a challenge. This is due to the complexity and wide range of interactions metals can perform. For instance, the strength of metal–ligand bonds is intermediate between that of non-bonded interactions and covalent bonds [28–30]. Moreover, in MD simulations of metal-binding proteins, variable coordination geometries, ligand-field and polarization effects add a layer of complexity [29–32]. Ultimately, accounting for transition metals in force fields is rather challenging [33]. Different classical modeling approaches for describing systems that include metal ions in MD simulations have been developed over the past years [34]. They include non-polarizable models, which treat electrostatic properties by fixed-point partial charges on atoms (such as bonded [35–37], non-bonded [38–40] and cationic dummy atom [41,42] models), and polarizable models, which use different strategies to explicitly take into account polarizability and changes in electronic charge distribution (such as fluctuating-charge [43,44], Drude oscillator [45–47] and induced-dipole [48,49] models).

In this study, we focus on the Nuclear Protein Localization Protein 4 (NPL4), a major cofactor protein of the AAA+ ATPase p97 (also known as valosin-containing protein or Cdc48 in yeast) [50–52]. p97 maintains the protein homeostasis of the cell and addresses proteotoxic stress by unfolding stable proteins to target them for subsequent proteasomal degradation [53,54]. NPL4, in complex with Ubiquitin Fusion Degradation 1 (UFD1) protein, binds p97 and directs it to proteasomal degradation pathways [55,56]. It has been shown that yeast Npl4, cooperatively with p97/Cdc48, interacts with the Lys48-linked polyubiquitin chains of the substrates targeted for degradation. Subsequently, ubiquitin and substrates are translocated and unfolded in p97/Cdc48 and then delivered to the proteasome for degradation [57,58].

NPL4 includes an N-terminal ubiquitin regulatory X-like (UBXL) domain, two zinc finger domains (ZF1 and ZF2), an Mpr1/Pad1 N-terminal (MPN) domain, and a C-terminal domain [59] (**Figure 1A-B**). In addition, mammalian NPL4 possesses an additional zinc finger domain at the C-terminus (NZF) [59–61] (**Figure 1A-B)**. The cryo-electron microscopy (cryo-EM) structures of *Chaetomium thermophilum* and *Saccharomyces cerevisiae* Npl4-Udf1-p97/Cdc48 complex [58,62,63] show that Npl4 anchors with both ZFs on top of the cis-side of p97/Cdc48 in a tower-shaped organization. Furthermore, single-particle cryo-EM of the human NPL4-p97 complex revealed that NPL4 assumes seesaw conformations due to the different interaction of ZFs with p97 before ATP hydrolysis, suggested to be important for the unfolding activity of p97 [59].

**Figure 1.**
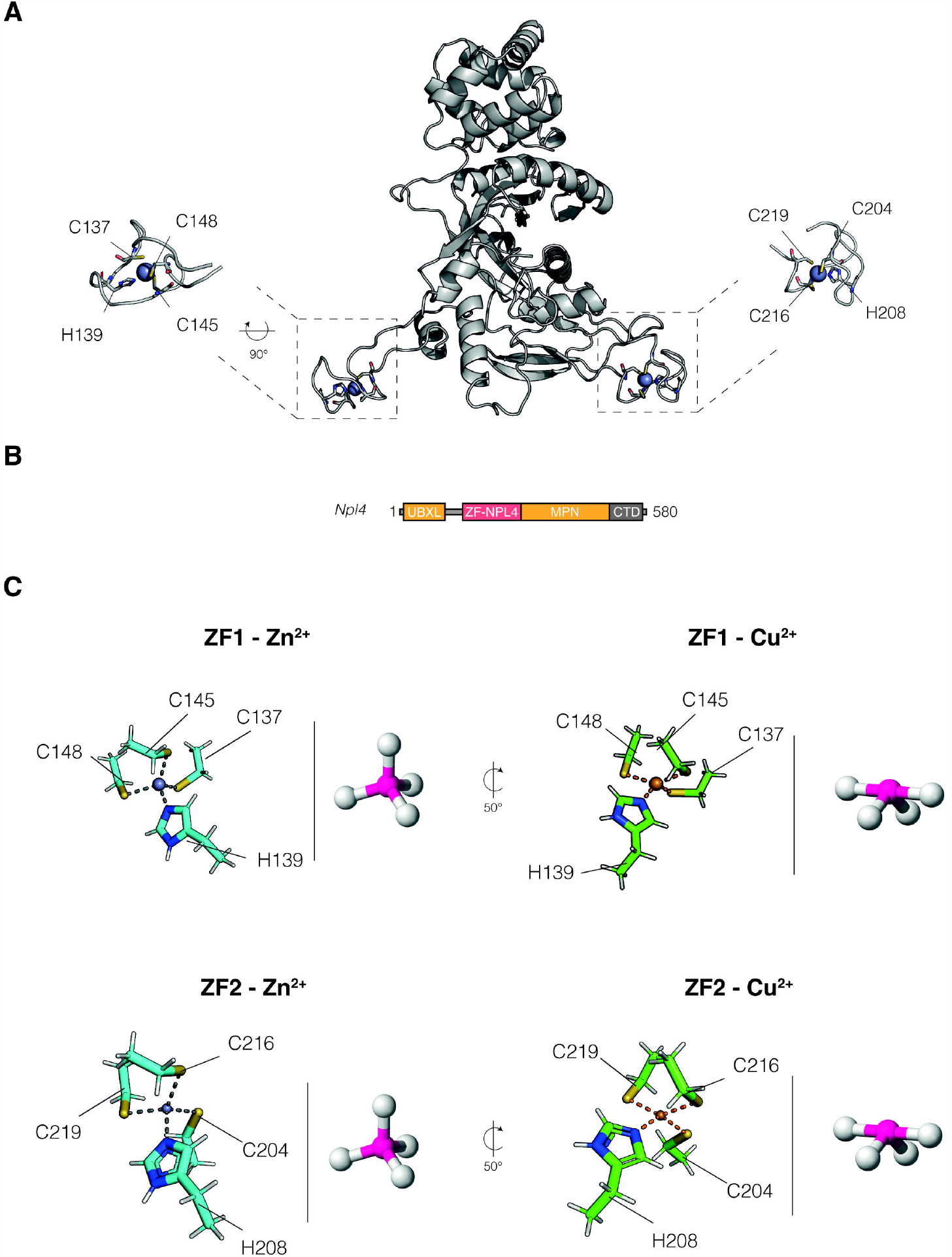
Overview of the system under study and expected coordination geometry. **(A)** We illustrate the yeast Npl4 structure used in the study. **(B)** Schematics diagram of yeast Npl4 protein domains. **(C)** Coordination geometry as estimated with QM calculations for geometry optimization for both ZF1 and ZF2 with Zn^2+^ and Cu^2+^ metals of the Npl4 protein. The obtained coordination geometries are compared to the tetrahedral and seesaw coordination. Images of the tetrahedral and seesaw conformation are from Wikimedia Commons.

It has been shown that the alcohol-abuse drug disulfiram (DSF), also known as Antabuse, targets and inhibits NPL4 in different cancer types [64]. However, we have only sparse knowledge about the molecular mechanisms of the anti-cancer activity of DSF. Fluorescence assay [65] and NMR data [66] showed that DSF acts as an ejector of zinc ions (Zn^2+^) from the zinc fingers of different proteins [67]. It has been suggested that DSF induces the ejection of Zn^2+^ from the zinc fingers by acting as an electrophile, mediating a thiol–disulfide exchange with the nucleophilic cysteine sulfurs [68,69]. Furthermore, recent evidence proposed that a metabolic derivative of DSF, bis-(diethyldithiocarbamate)-copper (CuET), plays a key role in targeting NPL4, inducing its unfolding and aggregation [64]. In addition, single-particle cryo-EM and in vitro and cell-based assays [59] suggested that cupric ions (Cu^2+^), released from CuET under oxidative stress, interact with ZF1 and ZF2 of NPL4, causing structural alterations and affecting the activity of NPL4. These effects could be related to the interaction and displacement of Zn^2+^ by Cu^2+^, as it is known that copper affects the structure and function of other zinc fingers [70–73].

In this study, we aim to provide, using classical MD simulations, a comparison between different force fields and their performances with different constructs of the yeast Npl4 protein. We started to examine non-bonded models because the unfolding mechanism of ZFs in NPL4 could rely on the detachment and replacement of Zn^2+^. We compared the results from MD simulations with optimized geometries from quantum mechanics (QM) calculations using model systems of the zinc coordination site for Npl4. We focused on one CHARMM force field, namely CHARMM36m [18], and two AMBER force fields, ff99SB-ILDN [74] and ff99SB^*^-ILDN [15,74]. Furthermore, we investigated the performance of an AMBER force field including bonded parameters for the Cu^2+^ site in Npl4 that we obtained based on QM calculations.

## 2. Materials and Methods

### 2.1 Quantum-mechanics calculations

The starting structures for QM calculations on ZF1 and ZF2 were extracted from the X-ray structure of yeast Npl4 (PDB ID: 6JWH, [63]). In the model systems, we kept only the atomic coordinates of Zn^2+^ and the metal-coordinating residues, i.e., C137–H139–C145–C148 and C204–H208–C216–C219 for ZF1 and ZF2 respectively (ZF1-Zn^2+^ and ZF2-Zn^2+^). We truncated the amino acids at the C_α_ positions and added a methyl group as a capping group (i.e. the ligands were modeled by CH_3_CH_2_S^−^ and ethyl imidazole, respectively). To study the effect of the possible displacement of Zn^2+^ by Cu^2+^, we also studied the same models with Zn^2+^ replaced by Cu^2+^ (ZF1-Cu^2+^ and ZF2-Cu^2+^). We performed geometry optimizations with Density Functional Theory (DFT) [75], first with the Cα coordinates frozen during calculations and then releasing them. On the final optimized geometries, we calculated harmonic vibrational forces. We calculated charges on all atoms with the restrained electrostatic potential (RESP) approach and using the Merz–Kollman scheme [76] with 10 concentric layers with density of points of 17 points/Å^2^). We also analyzed which coordination geometry Cu^2+^ assumes in models of ZF1 and ZF2 when in a coordination complex with only three residues (i.e., two cysteines and one histidine or three cysteines). We thus modeled the dissociation of one cysteine (C148 for ZF1 and C219 for ZF2) or one histidine (H139 for ZF1 and H208 for ZF2) by lengthening the distances between Cu^2+^ and the sulfur atoms or Nε2 nitrogen atom to 15 Å using GaussianView v6.1.1 [77]. We then performed geometry optimizations of the dissociated systems. For all the DFT calculations, we used the hybrid B3LYP exchange-correlation functional and the triple zeta TVZP basis set [75,78,79]. Furthermore, we used a polarizable continuum model with a dielectric constant of 4 to implicitly take into account the protein environment. All the input files and calculations were prepared and performed with GaussianView v6.1.1 and Gaussian v16, respectively [77,80].

### 2.2 Analysis of QM calculations

We used GaussianView v6.1.1 [77] to visualize the final optimized geometries and to calculate relevant geometry parameters of the four model systems, viz., atom–atom distances, angles, and dihedral angles. We also used this software to visualize the calculated vibrational modes. We used SHAPE v2.1 [81] to estimate the coordination geometry of the metal binding sites in the optimized model systems. SHAPE computes continuous shape measures (CShM) of the spatial coordinates of a set of points compared to vertices of ideal reference polygons or polyhedra, i.e., reference coordination geometries. Thus, we calculated CShM considering the coordinates of the atoms coordinating the metal ion (sulfur atoms for cysteines and nitrogen atoms for histidines). A CShM value equal to 0 corresponds to a structure matching the reference shape/coordination geometry perfectly. In contrast, the maximum allowed value is 100, corresponding to the hypothetical case where all vertices collapse into a single point in space. For example, in the case of the optimized geometry of ZF2-Zn^2+,^ we obtained CShM scores of 27.3 for planar square conformation, 1.2 for tetrahedral conformation, 6.1 for seesaw conformation, and 2.2 for vacant trigonal bipyramid conformation.

### 2.3 Development of bonded MD parameters

We used QM calculations to obtain a bonded model for AMBER force fields to treat Cu^2+^ and its binding atoms in the ligands (i.e., sulfur atom of cysteines and Nε2 nitrogen atom of histidines) in Npl4 (ff99SB^*^-ILDN-bonded). In our model, explicit bonds are defined between the metal and its ligands, which are treated as covalent bonds, i.e., including bond, angle, and dihedral terms. We used standard AMBER atom types for all metal-coordinating residues and a distinct atom type was used for Cu^2+^. Thus, we parametrized only the bonded interactions of Cu^2+^ and its coordinating atoms. All the other parameters were taken from ff99SB^*^-ILDN force field. We employed the Hess2FF program [83,84], based on the Seminario approach [82] to extract the force constants for bond, angles and dihedrals from the Hessian obtained from the QM frequency calculations. We used RESP charges derived from QM calculations for the atomic charges of the metal ion and the coordinating residues.

### 2.4 Structure selection for MD simulations

We collected all the available structures of Npl4 from the PDB and PDBredo databases (PDB IDs: 1Q5W, 6CHS, 6OA9, 6CDD, 6JWH, 6JWI, 6JWJ [58,61–63]). We evaluated the presence of missing atoms, alternative conformations using pdb4amber from AmberTools v20 suite [85]. For our MD simulations, we focused on two different systems: a) the zinc-finger domain ZF1 of the yeast Npl4 (Npl4(Zn^2+^)_129–151_, residues 129–151), and b) a construct without the UBXL N-terminal domain (residues 1–76) and the disordered linker (residues 77–112) of Npl4 (Npl4(Zn^2+^)_113–580_ (residues 113–580) (**Table 1**). We prepared the corresponding starting system in a complex with Cu^2+^ (Npl4(Cu^2+^)_113–580_) by replacing Zn^2+^ with Cu^2+^ and preserving its atomic coordinates. We employed the yeast structure of Npl4 (PDB ID: 6JWH [63]) to model the ZF1 Npl4(Zn^2+^)_129–151_ construct since it has a reasonable resolution (1.72 Å). Moreover, this structure represents the UFD1-unbound state. We used the original structure from the PDB database after verifying that the PDBREDO [86] refinement did not improve the quality of this structure. As mentioned above, we focused on the residues 129–151 to preserve the antiparallel β-bridge between P150–L151 and I129 and the mainchain hydrogen bond between I129 and L151. We used Modeller v10 and template structures with PDB IDs: 6JWI and 6JWJ [63] to remodel missing residues and retromutate back to the wild-type sequence mutations founds in the PDB file (E123A, K124A, and E125A) for the Npl4(Zn^2+^)_113–580_ construct. We did not include the N-terminal region 1–112 since none of the available yeast Npl4 structures in PDB database contain coordinates for this region that could be employed as templates. We defined the protonation of the histidines by visual inspection of the experimental structures of Npl4 and by predictions with PROPKA3 [87] and H++ [88]. In detail, we modeled the histidine residues 170, 205, 236, 290, 344, 468, 521, and 528 in the Nε2-H tautomeric state and 139, 159, 166, 208, 361, and 365 in the Nδ1-H tautomeric state. All aspartate, glutamate, arginine, and lysine residues were assumed to be charged. The cysteine ligands of the metal ions were assumed to be deprotonated, whereas all other cysteine residues were protonated.

**Table 1.** Summary of MD simulations and simulated systems.

| System | Force Field | Capping | Total atoms | Temperature (K°) | Length (Time) | N° replicates |
| --- | --- | --- | --- | --- | --- | --- |
| Npl4(Zn<sup>2+</sup>)<sub>129-151</sub> | CHARMM36m | NH2-CT2 | 17003 | 298 | 1 μs | 3 |
| Npl4(Zn<sup>2+</sup>)<sub>129-151</sub> | CHARMM36m | NH2-CT3 | 17003 | 298 | 1 μs | 1 |
| Npl4(Zn<sup>2+</sup>)<sub>129-151</sub> | ff99SB-ILDN | ACE-NH2 | 17332 | 298 | 1 μs | 3 |
| Npl4(Zn<sup>2+</sup>)<sub>129-151</sub> | ff99SB*-ILDN | ACE-NH2 | 17332 | 298 | 1 μs | 3 |
| Npl4(Zn<sup>2+</sup>)<sub>113-580</sub> | CHARMM36m | NH2-CT2 | 138909 | 298 | 500 ns | 3 |
| Npl4(Zn<sup>2+</sup>)<sub>113-580</sub> | ff99SB*-ILDN | ACE-NH2 | 139259 | 298 | 500 ns | 1 |
| Npl4(Cu<sup>2+</sup>)<sub>113-580</sub> | CHARMM36m | NH2-CT2 | 190986 | 298 | 500 ns | 3 |
| Npl4(Cu<sup>2+</sup>)<sub>113-580</sub> | ff99sb*-ILDN-bonded | NH2-CT2 | 139259 | 298 | 500 ns | 1 |

### 2.5 Molecular Dynamics (MD) simulations

The MD simulations were carried out with GROMACS v.2019 and they were run at a temperature of 298 K. We carried out three replicates of 1-μs all-atom explicit-solvent MD simulations for Npl4(Zn^2+^)_129–151_ with the CHARMM36m [18], ff99SB-ILDN [74], and f99SB^*^-ILDN [15,74] force fields (**Table 1**). We also performed three replicates of 500-ns MD simulations of Npl4(Zn^2+^)_113–580_ with CHARMM36m and one 500-ns simulation with ff99SB^*^-ILDN (**Table 1**). For the systems in which we replaced Zn^2+^ with Cu^2+^, we performed three 500-ns replicates for Npl4(Cu^2+^)_113–580_ with CHARMM36m. We capped the N- and C-termini of the starting models with either a hydrogen atom (NH2) and a –NH_2_ group (CT2) for the CHARMM force field, or with acetyl (ACE) and a –NH_2_ group (NH2) for the AMBER force fields. Furthermore, we collected an additional 1-μs simulations for Npl4(Zn^2+^)_129–151_ with CHARMM36m using a different C-terminal capping group (N-methyl amide, CT3 in **Table 1**). For the CHARMM simulations, we used parameters for Zn^2+^ [38], metal-coordinating residues [18], and Cu^2+^ [89] present in the CHARMM36m force field. For the AMBER force fields, we employed non-bonded parameters for cysteines and histidines for Zn^2+^-binding proteins [90]. The metals were modeled with a +2 charge and 12-6 Lennard-Jones potentials were used to model the Van der Waals interactions. We also collected a 500-ns MD simulation of Npl4(Cu^2+^)_113-580_ using the parametrized bonded model for Cu and the f99SB^*^-ILDN force field (ff99SB^*^-ILDN-bonded, **Table 1**).

As solvent models, we used TIP3P adjusted for the CHARMM36m force field (i.e., TIPS3P [91]) and TIP3P [92] for AMBER force fields. We solvated the systems in a dodecahedral box with a minimum distance of 20 Å between the protein and the edges of the box. We neutralized the charges of the systems using chlorine and sodium ions (between 2 and 19 ions depending on the system) and applied periodic boundary conditions. We equilibrated the systems in multiple steps. First, we performed a 10000-steps of minimization by the steepest descent method followed by 1 ns of solvent equilibration at 298 K, while restraining the coordinates of protein atoms by harmonic restraints with a force constant of 1000 kJ mol^−1^ nm^−2^. We then simulated the systems for 2 ns in the canonical ensemble (i.e., NVT) at 298 K using a *v*-rescale thermostat with a coupling constant of 6 ps. We then performed two simulations of 5 ns at a constant pressure of 1 bar (NPT ensemble). In the first NPT simulation, we used Berendsen barostat [93] with a coupling constant of 1 ps. We then selected a snapshot in the second half of the first NPT simulation in which the density of the system is close to the reference value (1.04 g/cm^3^). This frame was used as starting system for the second NPT simulation in which we used the Parrinello–Rahman barostat [94,95] with a coupling constant of 1 ps. Using the same approach, we selected a snapshot in the second half of the second NPT simulation as starting system for additional 5 ns in the NVT ensemble of 298 K using the *v*-rescale thermostat.

We performed the production MD simulations in the canonical ensemble at 298 K using a time-step of 2 fs. All covalent bonds were constrained with the LINCS algorithm [96]. We used the particle-mesh Ewald summation scheme to calculate long-range electrostatic interactions with a grid spacing of 1.2 Å and a cutoff of 9 Å for both Van der Waals and Coulomb interactions [97,98].

### 2.6 Analysis of MD simulations

We verified that the distance between the protein and its periodic images was always higher than the cutoffs used for the electrostatics or Van der Waals interactions (distance in the range of 65 ± 5 Å in our simulations) to avoid artifacts due to contact between the protein and one of its images [99]. We calculated the distances between the metal ion and the sulfur atoms or ε-nitrogen atoms of the side chains of the coordinating residues (C_137_–H_139_–C_145_–C_148_ and C_204_–H_208_–C_216_–C_219_ for ZF1 and ZF2 of Npl4) along the simulation time. We analyzed both the time series and the distribution.

We used the software SHAPE v2.1 [81] to estimate the coordination geometry of the metal binding sites in the simulations. To consider the impact of water molecules on the coordination geometry of the two metals in the CHARMMM36m replicates, we computed CShM including the oxygen atoms of water molecules.

We used the software CONtact Analysis (CONAN) [100] to analyze inter-residue and metal–residue contacts along the MD simulations. We used a cutoff *rcut* value of 10 Å, *rinter* and *rhigh-inter* values of 5 Å, using a protocol employed in a previous study [101]. We estimated the occurrence (i.e., the ratio between the number of frames in which a contact is present and the total number of frames of the simulations) and the number of encounters (i.e., how many times the contact is formed and broken) of each contact. As a reference, we used CONAN with the same parameters to analyze inter-residue and metal–residue contacts in the starting structure of Npl4(Zn^2+^)_129-151_.

## 3. Results

### 3.1. Coordination geometry in the zinc-finger binding sites of Npl4

First, we examined by DFT calculations model systems of the zinc coordination sites for yeast Npl4, including only the atomic coordinates of Zn^2+^ and the metal-coordinating residues for ZF1 and ZF2 (ZF1-Zn^2+^ and ZF2-Zn^2+^) (**Figure 1C**). To study the effect of the possible displacement of Zn^2+^ by Cu^2+^, we also performed calculations in which we replaced Zn^2+^ with Cu^2+^ in the initial structures (ZF1-Cu^2+^ and ZF2-Cu^2+^) (see Methods). We performed geometry optimization calculations at the B3LYP/TZVP level of the four model systems. The DFT-optimized geometries of the complexes are shown in **Figure 1C**, and we calculated different geometry parameters for the four metal-coordinating residues (**Figure 1C and Table 2**). We performed vibrational frequency calculations for the optimized geometry of each system, which showed that all frequencies have positive values, confirming that we found local minima.

**Table 2.** Geometric parameters from the QM geometry optimizations. The experimental X-ray structure used as a reference is 6JWH. ‘QM’ indicates the values estimated from the final structures after geometry optimization using DFT. The atoms in the table below are defined as following : M= Zn^2+^ / Cu^2+^; S1= C137(Sγ) / C204(Sγ); N1= H139(Nε) / H208(Nε); S2= C145(Sγ) / C216(Sγ); S3= C148(Sγ) / C219(Sγ); Cα1=C137(Cα)/C204(Cα); Cα2= H139(Cα) / H208(Cα); Cα3= C145(Cα) / C216(Cα); Cα4= C148(Cα) / C219(Cα); Cβ1= C137(Cß) / C204(Cß); Cβ2= H139(Cß) / H208(Cß); Cδ2= H139(Cδ2) / H208(Cδ2); Cβ3= C145(Cß) / C216(Cß); Cβ4= C148(Cß) / C219(Cß).

| System | ZF1-Zn<sup>2+</sup> | ZF1-Zn<sup>2+</sup> | ZF1-Cu<sup>2+</sup> | ZF2-Zn<sup>2+</sup> | ZF2-Zn<sup>2+</sup> | ZF2-Cu<sup>2+</sup> |
| --- | --- | --- | --- | --- | --- | --- |
|  | X-ray | QM | QM | X-ray | QM | QM |
| **CShM score** |  |  |  |  |  |  |
| Square (SP-4) | 27.76 | 28.51 | 12.56 | 32.88 | 27.28 | 14.46 |
| Tetrahedron (T-4) | 0.81 | 1.17 | 6.85 | 0.44 | 1.22 | 5.69 |
| Seesaw (SS-4) | 6.70 | 6.26 | 3.83 | 8.53 | 6.08 | 3.90 |
| Vacant trigonal bipyramid (vTBPY-4) | 2.43 | 1.56 | 8.22 | 1.87 | 2.16 | 7.39 |
| **Distances (Å)** |  |  |  |  |  |  |
| M-S1 | 2.31 | 2.38 | 2.34 | 2.35 | 2.38 | 2.34 |
| M-N1 | 2.09 | 2.16 | 2.12 | 2.09 | 2.15 | 2.13 |
| M-S2 | 2.34 | 2.38 | 2.34 | 2.31 | 2.38 | 2.35 |
| MS3 | 2.31 | 2.39 | 2.35 | 2.27 | 2.39 | 2.33 |
| **Angles (°)** |  |  |  |  |  |  |
| M-S1-Cβ1 | 98.87 | 106.85 | 111.49 | 99.02 | 104.75 | 108.95 |
| M-N1-Cδ2 | 133.19 | 127.57 | 125.21 | 131.43 | 127.22 | 125.07 |
| M-S2-Cβ3 | 111.85 | 104.81 | 106.02 | 109.2 | 105.58 | 106.73 |
| M-S3-Cβ4 | 98.29 | 103.98 | 108.55 | 103.87 | 103.63 | 107.28 |
| S1-M-S2 | 107.65 | 113.2 | 100.14 | 111.55 | 104.86 | 92.88 |
| S1-M-S3 | 109.57 | 108.98 | 96.96 | 112.13 | 113.41 | 105.03 |
| S1-M-N1 | 113.51 | 108.28 | 132.92 | 105.67 | 111.45 | 130.29 |
| S2-M-N1 | 106.24 | 103.36 | 98.45 | 106.96 | 104.76 | 100.15 |
| S2-M-S3 | 119.58 | 122.63 | 144.48 | 114.55 | 123.13 | 141.21 |
| S3-M-N1 | 100.26 | 98.22 | 92.16 | 105.19 | 98.59 | 94.00 |
| **Dihedral (°)** |  |  |  |  |  |  |
| Cα1-Cβ1-S1-M | -95.47 | -85.03 | -100.67 | -142.24 | -88.39 | -176.28 |
| Cα2-Cβ2-N1-M | -60.48 | -98.02 | -103.39 | -96.03 | -104.6 | -100.6 |
| Cα3-Cβ3-S2-M | -73.46 | -93.93 | -73.76 | -74.57 | -88.55 | -74.58 |
| Cα4-Cβ4-S3-M | -108.53 | -78.6 | -69.66 | -97.99 | -79.55 | -68.59 |

With respect to the experimental structure of ZF1, the optimized structure of ZF1-Zn^2+^ did not present any major rearrangements, and the tetrahedral coordination of the system was preserved, with only minor rearrangements for some dihedral angles (**Table 2**). We used SHAPE to estimate the coordination geometry (**Figure 1C and Table 2**). For ZF1-Zn^2+^ and ZF2-Zn^2+^, we observed that the lowest CShM scores are associated with a tetrahedral geometry (1.2). These results agree with known properties and coordination geometries of Zn^2+^ in zinc finger domains [102–104].

In contrast, the optimized structures of ZF1/ZF2-Cu^2+^ showed significant structural rearrangements (**Figure 1C and Table 2**). In particular, the angles between the sulfur of the thiol group of C137 and the metal and the nitrogen of the imidazole ring of H139 in ZF1 or the sulfur atoms between C216 and C219 in ZF2 increase (from 109° to 133° and from 123° to 141°). Cupric ions generally prefer tetragonal structures, possibly with weaker axial ligands [105], but this can change with sulfur ligands. We observed the lowest CShM score for the seesaw conformation (3.8 and 3.9 for ZF1 and ZF2), respectively, suggesting that ZFs-Cu^2+^ systems adopted a distorted seesaw conformation (**Figure 1C and Table 2**). We investigated if, in our systems, Cu^2+^ has the propensity to switch to a trigonal conformation by dissociation of one of the coordinating groups. We thus modeled on the optimized structure of ZF1-Cu^2+^ and ZF2-Cu^2+^ the dissociation of one cysteine or one histidine by lengthening their Cu^2+^-distance to 15 Å. After geometry optimization by DFT calculations, we observed that the four dissociated systems assume a trigonal planar conformation (CShM scores between 1.1–1.8). Comparing the minimum energies calculated for the dissociated and the corresponding four-coordinate systems, we noticed that the seesaw conformations are the most stable for ZF1 and ZF2 in the presence of Cu^2+^.

In the following, we will use the QM-derived geometries as a reference compared to the coordination observed during the MD simulations.

### 3.2 The zinc-finger domain alone is not a good model system to study Npl4 in MD simulations

First, we evaluated if we could use the construct Npl4(Zn^2+^)_129-151_ as a minimalistic model system to study the effects of the replacement of Zn^2+^ with Cu^2+^ on NPL4 by MD. Npl4(Zn^2+^)_129-151_ includes ZF1 of Npl4 only. To this goal, we run 1-μs unbiased MD-simulation at 298 K with four combinations of force fields and capping modes (see **Methods, Table 1**; all using a non-bonded model for Zn^2+^).

First, we evaluated how the various force fields describe the distances between the zinc ion and its four coordinating residues (i.e., C137, H139, C145, C148) (**Figure S1)**. We observed that most force fields featured minor deviations from the reference QM distances (< 0.2 Å). The main exception was CHARMM36m. In particular, most of the CHARMM36m replicates featured stable coordination distances between the four residues and the metal ion (**Figure 2A, Figure S1**). However, in replicate 2, Cys148 dissociated from the Zn^2+^ ion after 350 ns (**Figure S2**). In addition, we observed one short detachment event in replicate 1, at around 300 ns, between C148 and the Zn^2+^ ion. This brief event lasted approximately 30 ps and during which the distance increased up to 4.5 Å. To better evaluate the timing of these events and the structural consequences, we used a contact-based method to evaluate the occurrence of interactions between the Zn^2+^ ion and the protein residues (**Figure 2B**). In the X-ray structure used for the simulation, this method identified K133, S134, K138, M144, Y147, and the coordinating residues within 5 Å of distance as a reference. In the case of replicate 2, C148 was found at lower occurrence (> 0.5) and with a few encounter events in line with the fact that we do not observe any reassociation after the dissociation of C148. Also, K133 and S134 make contacts with lower occurrence than in the other replicates, where we observed stable metal coordination. In addition, Q132 and L136 populate the surroundings of Zn^2+^ when the cysteine detaches. To further investigate the detachment of C148 from Zn^2+^ in replicate 2, we analyzed the occurrence of atomic contacts of C148 before and after the detachment (**Figure S3**). In the starting structure, we identified Zn^2+^ and residues R131-S134, C137, H139, and G143-P150 in the vicinity of C148. We observed that after the detachment, the surrounding of C148 is not preserved, showing a lower occurrence of residues belonging to the coordination center, i.e., H139 and C145, and instead starts having transient contacts with other residues in the N- and C-terminal regions, such as I129-L136 (**Figure S3**). We observed that the sulfur atom in the side chain of C148 is close to the backbone amide of C145 (around 3.3 Å). It has been suggested that hydrogen bonds between the sulfur atom of cysteines and backbone amides have a role in the reactivity of zinc fingers [68]. Furthermore, we observed that the detachment of C148 causes the system to sample unfolded and distorted conformations of Npl4(Zn^2+^)_129–151_.

**Figure 2.**
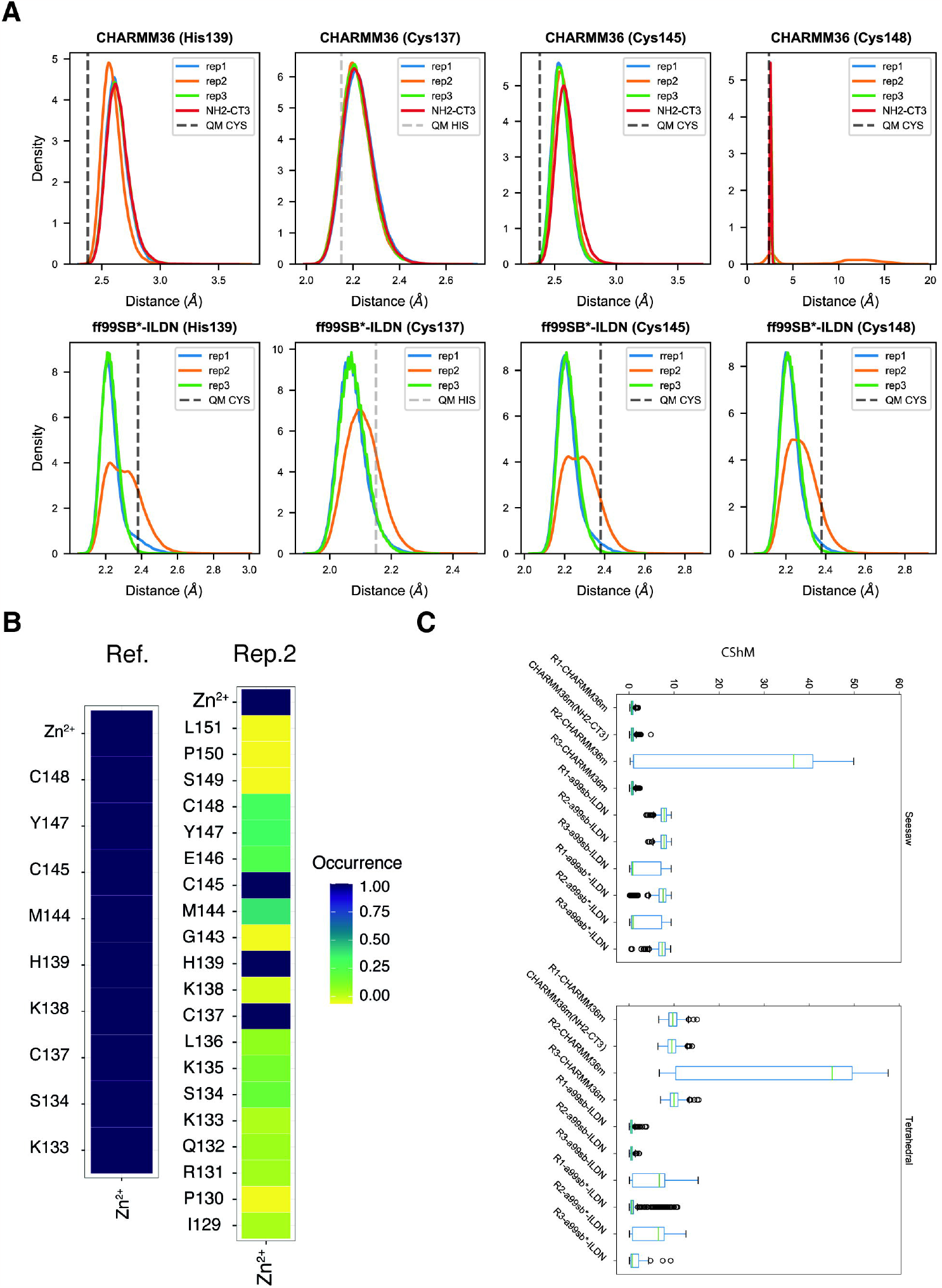
Coordination distances and geometries in Npl4(Zn^2+^)_129-15_. **(A)** Distance distributions of all metal-coordinating residues for Npl4(Zn^2+^)_129-151_ CHARMM36m and ff99SB^*^-ILDN replicates. In dashed lines are the reference values from QM calculations. The reference values for the cysteines were obtained by averaging the distances obtained in the QM calculations. **(B)** Occurrences of contacts for Zn^2+^ ion between Npl4(Zn^2+^)_129-151_ CHARMM36m replicate two and reference starting structure. The label “Ref.” denotes the reference structure, while “R2” replicate two. **(C)** Boxplot of CShM scores grouped by tetrahedral and seesaw conformations for all Npl4(Zn^2+^)_129–151_ MD replicates. R1, R2, R3 stand for replicate 1, replicate 2 and replicate 3.

Next, we evaluated the coordination geometry for each simulation of Npl4(Zn^2+^)_129–151_ and compared them to those estimated by the DFT calculations (**Table S1, Figure 2C**). We used SHAPE to estimate the coordination geometry of the metal ion in the various replicates of Npl4(Zn^2+^)_129–151_. The analysis showed that replicates where detachment happens generally deviated from all the reference four-coordinate geometries (showing average CShM scores > 20). For the AMBER force fields, we observed stable coordination distances between the four residues, which preserved the tetrahedral coordination geometry for replicates 1–2 of ff99SB-ILDN and replicates 1 and 3 of f99SB^*^-ILDN (average CShM scores 0.6–1.5, **Table S1, Figure 2C**), in agreement with the QM-derived references. On the other hand, the other two replicates were closer to seesaw conformations. We observed similar behavior with CHARMM36m, noting that replicates with Zn^2+^ preferred seesaw conformations (average CShM scores between 0.7–0.8), in disagreement with the results obtained by the QM calculations (**Table S1, Figure 2C**).

Therefore, we investigated the behavior of water molecules around the Zn^2+^ ion during the MD simulations. We observed that in all the CHARMM36m replicates, two water molecules coordinate Zn^2+^ during all the trajectories (average distance 2.2 Å), increasing the coordination number of Zn^2+^ to six. Including the two Zn^2+^-coordinating water molecules in the SHAPE analysis, we noticed that the CHARMM36m replicates 1 and 3 (replicate 2 undergoes detachment events) result in octahedral coordination geometries (average CShM scores 1.6, **Table S1**). On the other hand, for the AMBER force fields, we did not observe any additional water molecules in the coordination shell of Zn^2+^ nor any tendency to increase the coordination number (**Table S1**).

These results indicate that the force fields for which we have used the non-bonded parameters available for GROMACS cannot provide any stable and reliable microsecond simulations of the Zn^2+^ ion in Npl4(Zn^2+^)_129-151_, and this system cannot be used as a minimal model for our study. This is a mandatory requirement before using the simulation protocol to understand the structural changes that Cu^2+^ could cause on the Npl4 structure.

### 3.3 Zn^2+^ and Cu^2+^ coordination in the context of the full-length protein

In light of the results described above, we turned our attention to a larger construct (residues 113–580, Npl4(Zn^2+^)_113-580_). We aimed to evaluate if the simulation of the system accounting for the full structural environment around the ZF1 binding site could compensate for the low stability of the coordination site during the MD simulations of the zinc finger domain construct. These simulations also allowed us to monitor the behavior of ZF2. We evaluated how the different force fields described the distances between the zinc ion and its four coordinating residues in Npl4(Zn^2+^)_113-580_ for ZF1 (i.e., C137, H139, C145, and C148) and ZF2 (i.e., C204, H208, C216, and C219). We observed that all force fields featured minor deviations from the reference QM distances (< 0.4 Å), showing that they preserved stable coordination with Zn^2+^for both ZF1 and ZF2 (**Figure S4**). In addition, we assessed the coordination geometry for each simulation of Npl4(Zn^2+^)_113–580_ using SHAPE (**Figure 3**). We found that the ff99SB^*^-ILDN force field had a preference for a tetrahedral conformation for ZF1 over a vacant trigonal bipyramidal conformation, while ZF2 had a very slight preference for a vacant trigonal bipyramid conformation (Figure 3, **Table S1**). On the other hand, all CHARMM36m replicates preferred a seesaw conformation for both ZFs, in contrast to the QM calculations (the CShM scores for replicates 1–3 are 0.8–1.2, **Table S1**). The time series of the CShM scores (**Figure S5**) also confirm the assignments in **Table S1** and showed that in the ff99SB^*^-ILDN simulation while ZF1 preserved a tetrahedral conformation, ZF2 oscillated between tetrahedral, vacant trigonal bipyramidal and seesaw conformations.

**Figure 3.**
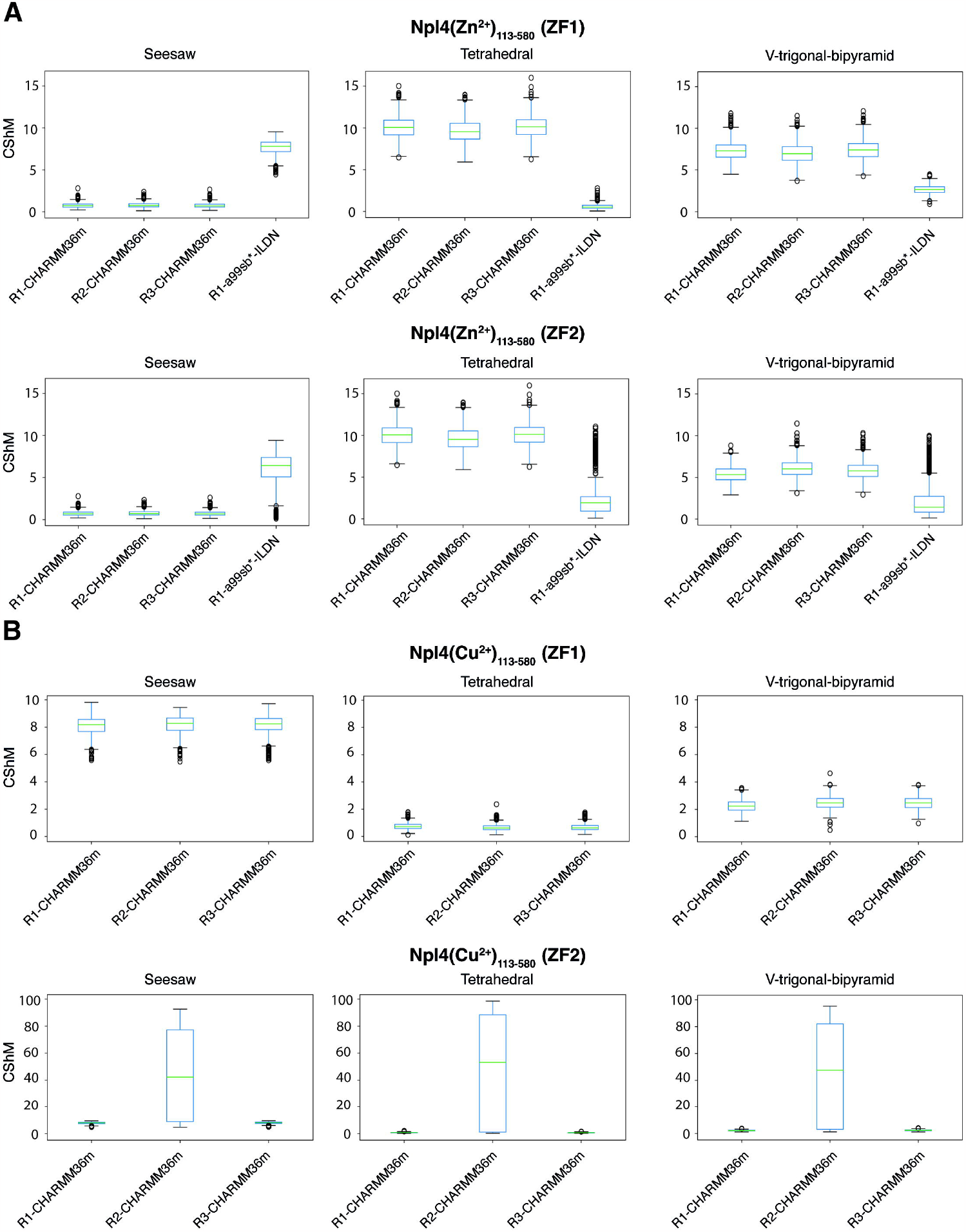
Coordination distances and geometry in Npl4(Zn^2+^)_113-580_ and Npl4(Cu^2+^)_113-580_. **(A)** Boxplot of CShM scores grouped by tetrahedral, seesaw and vacant trigonal bipyramid conformations and divided by ZFs domain of all Npl4(Zn^2+^)_113-580_ MD replicates. R1, R2, and R3 stand for replicate 1, replicate 2, replicate 3. **(B)** Similar boxplot to (A) for Npl4(Cu^2+^)_113–580_. Noticeable is the high CShM score for ZF2 in replicate two, where the dissociation of the cupric ion happens.

We noticed that the CHARMM36m simulations featured two stable water molecules in the Zn^2+^ coordination site for ZF1, as well as one water molecule and the backbone oxygen of C204 for ZF2 (the average distance is around 2.1 Å) (**Figure 4A)**. This is the reason why we get a seesaw conformation for the zinc-bound form of Npl4. Therefore, we also included the water molecules in the SHAPE calculation to analyze the hexa-coordination of Zn^2+^ in ZF1 and ZF2 (**Figure 4B and Table S1**). We observed that all CHARMM36m replicates adopted octahedral coordination geometries for ZF1 and ZF2 (the average CShM scores are between 1.5 and 1.9–2.0, respectively; **Table S1**). As observed for the minimal systems Npl4(Zn^2+^)_129-151_, the non-bonded model in CHARMM36m showed a strong tendency to favor the entrance of additional water molecules in the coordination shell of Zn^2+^, increasing the coordination number to six and adopting octahedral coordination geometries. Octahedral geometries of Zn^2+^ with six water molecules dominate in aqueous solution [106,107]. Thus, the performance of non-bonded parameters on our model systems using CHARMM36m reflects how the parameters have been developed, referring to complexes of Zn^2+^-water molecules [38]. On the other hand, for the ff99SB^*^-ILDN force field, we did not observe any additional water molecules in the coordination shell of the metal.

**Figure 4.**
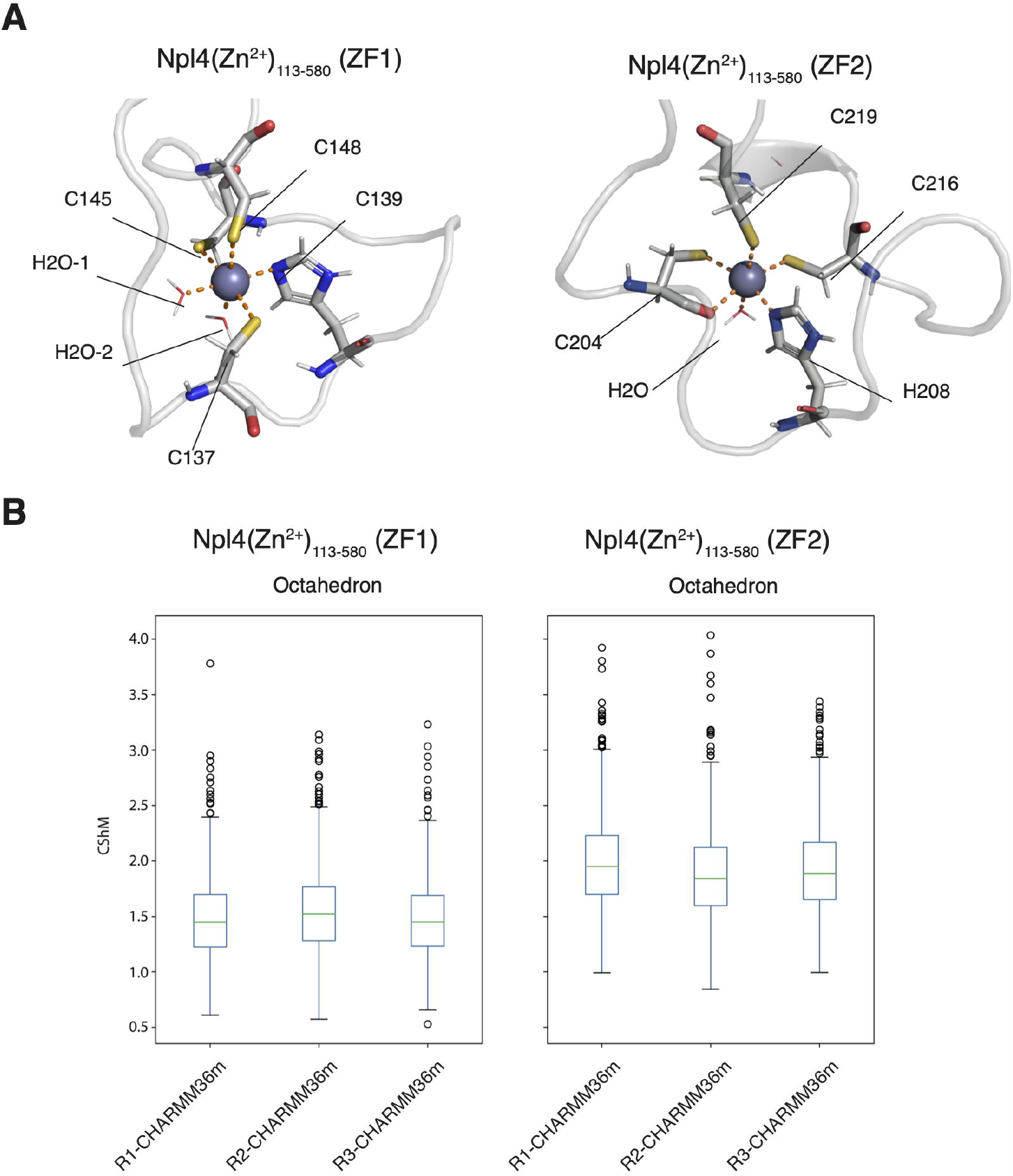
Interaction with water in the metal binding site of Npl4(Zn^2+^)_113–580_. **(A)** Snapshot of ZF1 and ZF2 of Npl4(Zn^2+^)_113–580_ from CHARMM36m replicate one, which highlights the zinc fingers coordination by water molecules. In the case of ZF2, the oxygen atom of C204 backbone coordinates the metal ion instead of a second water molecule. **(B)** Boxplot of CShM scores of all Npl4(Zn^2+^)_113–580_ CHARMM36m replicates grouped by octahedral configuration and divided by ZFs domain. R1, R2, and R3 denote replicate 1, replicate 2, and replicate 3, respectively.

A disagreement with the QM calculations was also observed when investigating the coordination geometry with the Cu^2+^ ion in the CHARMM36m simulations of Npl4(Zn^2+^)_113–58_. Indeed, all replicates adopted a tetrahedral conformation for ZF1 and ZF2 (except for ZF2 in replicate 2, in which the Cu^2+^ ion dissociated). This behavior was also confirmed by the time series of CShM scores (**Figure 3B and S5**). On the other hand, we did not observe any additional water molecules in the coordination shell of the Cu^2+^ ion.

The 12-6 Lennard-Jones parameters employed for Cu^2+^ in CHARMM36m have been parametrized in metal-water complexes using free energy perturbation simulations to reproduce experimental free energies of hydration [89]. The parameters for Zn^2+^ and coordinated cysteines and histidines used for ff99SB^*^-ILDN have recently been developed and validated based on QM calculations by redetermining the atomic charges of the coordinating atoms of the residues and adjusting non-bonded Lennard-Jones parameters [90]. Our simulations confirmed the overall improved performance of these parameters in the AMBER force field, which better preserved the tetra-coordination of the Zn^2+^ binding site in Npl4(Zn^2+^)_113–580_ (**Table S1**).

### 3.4 Bonded model of Cu^2+^ in the zinc-finger binding sites of Npl4

As a results of dissociation events and distortion of the coordination geometries in the MD simulations with non-bonded models, we decided to also evaluate the effectiveness of a bonded model. We obtained a bonded model for AMBER force fields to describe Cu^2+^ and its binding atoms in the ligands in Npl4. We performed a parametrization of the Cu^2+^ coordination site from our QM calculations using the Hess2FF implementation of the Seminario method [82]. Then, we carried out a 500-ns MD simulation of Npl4(Cu^2+^)_113–580_ using the calculated bonded parameters and f99SB^*^-ILDN force field (ff99SB^*^-ILDN-bonded, in **Table 1**).

We evaluated how the bonded model described the distances between Cu^2+^ and its four coordinating residues in Npl4(Cu^2+^)_113–580_ for ZF1 and ZF2 (**Figure 5A**). We observed that ff99SB^*^-ILDN-bonded preserved the coordination geometries of ZF1 and ZF2 through all the simulation. As expected, the bonded model performed better than the tested non-bonded models in obtaining the correct metal–ligand bond lengths, showing almost identical distances compared to the reference QM values. We observed that water molecules occasionally come close to the Cu^2+^ ion (at a distance minor or equal to 3.5 Å for around the 5 and 10 % of the simulation frames for ZF1 and ZF2, respectively) but we did not observe any water molecules stably in the coordination shell of the metal.

**Figure 5.**
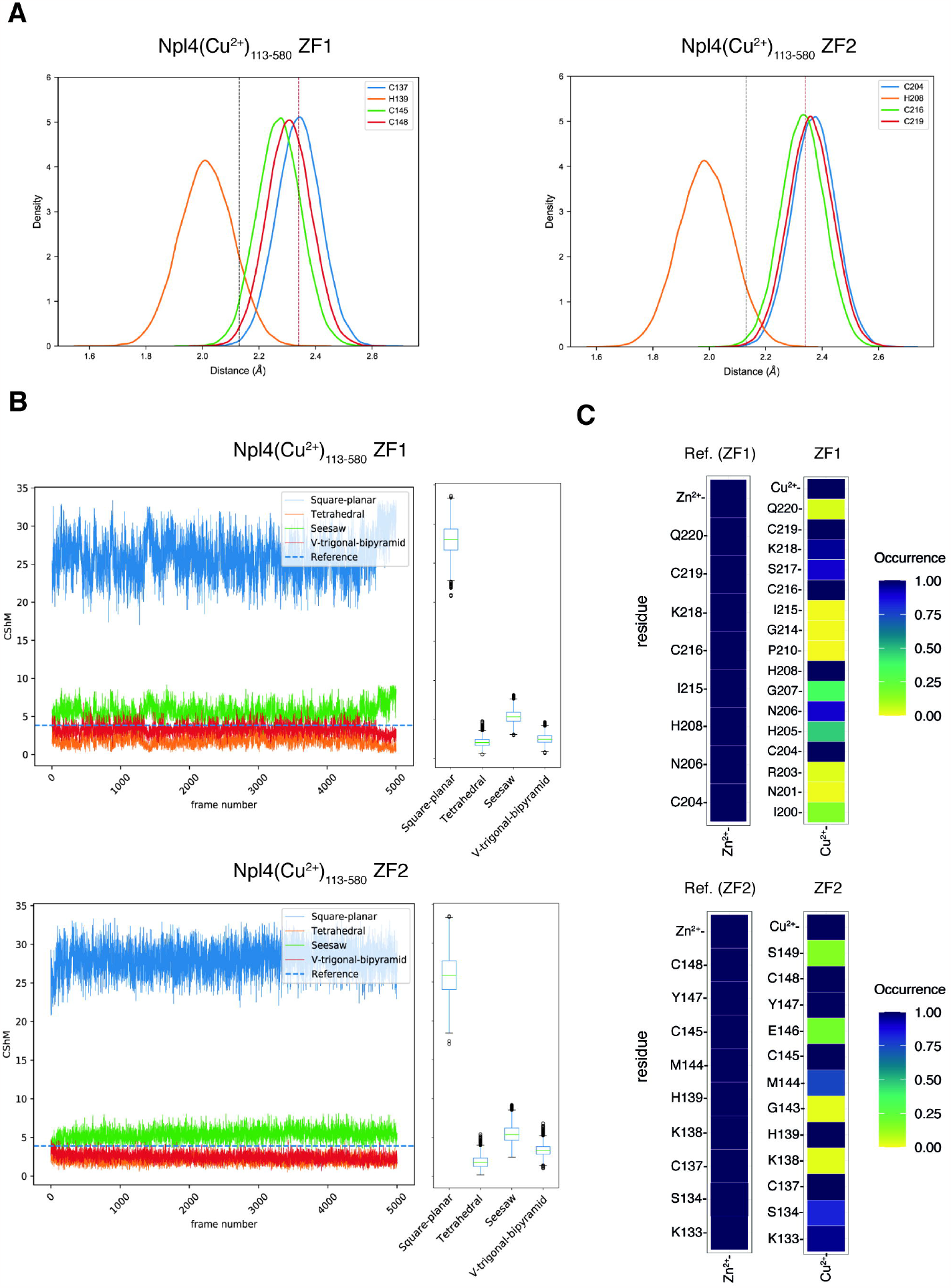
Coordination distances and geometry for the bonded model of Npl4(Cu^2+^)_113–580_ ff99sb^*^-ILDN. **(A)** Distance distributions of all metal-coordinating residues for ZF1 and ZF2 of Npl4(Cu^2+^)_113–580_ ff99sb^*^-ILDN with bonded model. The dashed lines are the reference values calculated for the QM optimized structure of ZF1-Cu^2+^ and ZF2-Cu^2+^. The black lines indicate the values calculated for the histidines, while the brown lines indicate reference values calculated for the cysteines. For the latter, the values were obtained by averaging the distances obtained in the QM calculations for the three cysteines in ZF1 and ZF2. **(B)** Time series and boxplots of CShM scores of ZF1 and ZF2 of Npl4(Cu^2+^)_113–580_. The dotted lines indicate the CshM score for the seesaw conformation of the QM optimized structure of ZF1-Cu^2+^ and ZF2-Cu^2+^. **(C)** Occurrences of contacts between Cu^2+^ and surrounding residues in the ZF1 and ZF2 of Npl4(Cu^2+^)_113–580_ ff99sb^*^-ILDN with bonded model and reference starting model. The label “Ref.” denotes the reference structure.

We also assessed the coordination geometry of Npl4(Cu^2+^)_113-580_ for ZF1 and ZF2 using SHAPE (**Figure 5B**) and evaluated the occurrence of interactions between the Cu^2+^ ion and the protein residues (**Figure 5C**). We observed that both ZF1 and ZF2 oscillated between tetrahedral, vacant trigonal bipyramidal and seesaw conformations, with a slight preference for tetrahedral and vacant trigonal bipyramid conformations (the average CShM scores are 1.9–2.1 and 2.5–3.4, respectively; **Table S1**) over seesaw conformations (the average CShM scores are between 5.4–5.5 **Table S1**).

Compared to the starting structure, the surrounding environment of the metal ion is largely preserved during the MD simulation for both ZF1 and ZF2 (**Figure 5C**). The contact analysis identified the metal-coordinating residues (C137, H139, C145, C148 for ZF1 and C204, H208, C216, C219) and K133, S134, K138, M144, Y147 for ZF1 or N206, S217, K218 for ZF2 with high-occurrence within 5 Å of distance of the metal ion. Overall, the results obtained with ff99SB^*^-ILDN-bonded reproduce the QM calculations for the coordination geometry of Cu^2+^ ion in which we observed a distorted seesaw conformation. In future, our parametrization could be improved by employing more complex and accurate methods, including the definition of special atom types and parameters for each metal ligand.

## 4. Discussion

The alcohol-abuse drug Antabuse has recently been repurposed for cancer treatment by targeting and inhibiting NPL4 [64], a zinc-binding protein that contributes to regulating protein homeostasis as a cofactor for p97 [50–52]. Recent experimental studies proposed that Antabuse metabolites and cupric ions released from them induce NPL4 misfolding and aggregation [59,64]. In this study, we aimed to evaluate force-field parameters for metal ions for MD simulations could reliably describe the structure and dynamics of NPL4.

Molecular simulations are becoming indispensable in studying structure-function-dynamic relationships of proteins: an open challenge is to model metal ions and metal-coordinating proteins accurately [26,27,34]. However, accurate classical modeling of zinc-coordinating proteins, such as NPL4, is challenging. For example, current MD force fields have difficulties in accurately describing the chemical properties and complicated chemical bonding of zinc, such as its versatility to bind a wide range of ligands, including nitrogen, oxygen, and sulfur atoms, and its flexibility in forming coordination complexes with multiple coordination numbers. Furthermore, classical MD force fields lack an accurate description of charge transfer and polarization effects, which could be highly relevant in describing metal coordination complexes in proteins. Thus, a pivotal step to apply MD simulations to NPL4 and study the interactions with cupric ions and Antabuse metabolites and the consequent misfolding mechanism is first to identify a suitable force field to describe the protein in its zinc-bound state. Since a possible step in the misfolding mechanism of ZFs in NPL4 could involve detachment and replacement of Zn^2+^ with Cu^2+^ ions, we focused on investigating the performance of non-bonded (12-6 Lennard-Jones) MD force fields to describe Zn^2+^ and Cu^2+^ ions and their interactions.

In our work, we provide a comparison between different Zn^2+^ force fields (CHARMM36m [18], ff99SB-ILDN [74] and f99SB^*^-ILDN [15,74]) and their performances in studying two constructs of *S. cerevisiae* Npl4 protein: a minimal model system including only the ZF1 motif of Npl4 (Npl4(Zn^2+^)_129–151_) and a larger construct including ZF1 and ZF2 motifs of Npl4 (Npl4(Zn^2+^)_113–580_). We then investigated the performance of the CHARMM36m non-bonded parameters for Cu^2+^ in Npl4(Cu^2+^)_113–580_, which may be involved in the possible mechanism by which copper replaces zinc in the ZF motifs. We focused on the ability of the force fields to model the coordination geometry of the metal ions by comparing the results from the MD simulations with optimized geometries from QM calculations at the DFT level of model systems of the Zn^2+^ and Cu^2+^ coordination site of ZF1 and ZF2 of Npl4.

Our data confirmed the importance of carefully designing and testing the protein construct and simulation protocol when studying metal coordination sites. We observed that the force fields could not provide microsecond stable coordination trajectories of the Zn^2+^ ion in Npl4(Zn^2+^)_129–151_. Instead, we identified detachment events of metal-coordinating residues, such as C148, and sampling of unfolded and distorted conformations. Overall, these limitations question the use of Npl4(Zn^2+^)_129-151_ as a minimal model system to study structural changes associated with metal replacement. On the other hand, for the simulations of the larger construct Npl4(Zn^2+^)_113-580_ we observed stable coordination of Zn^2+^ for both ZF1 and ZF2. Our results suggest that considering the full structural environment around the ZF1 coordination site compensates for the low stability observed during the MD simulations of Npl4(Zn^2+^)_129–151_.

Furthermore, our results showed that the combination of force fields and non-bonded parameters tested on Npl4(Zn^2+^)_113-580_ can generally provide reasonable descriptions of Zn^2+^ and Cu^2+^ coordination distances with cysteines and histidines compared to QM data. However, our simulations showed that non-bonded models have problems to maintain the proper coordination number and tetrahedral coordination geometry of Zn^2+^ in ZF1 and ZF2 of Npl4. We observed that non-bonded parameters for Zn^2+^ in CHARMM36m had a strong tendency to increase the metal coordination number to six and assumed octahedral coordination geometries by including additional water molecules in the coordination shell of the metal. The performance of these parameters in our model systems reflected how they had been developed using as reference the complexes of Zn^2+^-water molecules [38]. On the other hand, we observed that the parameters for Zn^2+^ and coordinated cysteines and histidines used for ff99SB^*^-ILDN, which have been recently parametrized based on QM calculations [90], gave an improved performance, better preserving the tetrahedral coordination of the Zn^2+^ binding site. For the non-bonded parameters for Cu^2+^ in CHARMM36m [89], we observed a preference to assume tetrahedral coordination geometries instead of the seesaw conformation suggested by our QM calculations. Overall, these discrepancies with the QM calculations did not allow us to use the non-bonded parameters in CHARMM36m for understanding the structural changes that Cu^2+^ could cause on the Npl4 structure. This is quite natural as a non-bonded model is spherically symmetric and therefore will never be able to model ligand-field effects that are important for Cu^2+^ (but not for Zn^2+^).

Therefore, we instead developed a bonded model for Cu^2+^, based on QM frequency calculations. Simulations with this model was more accurate in preserving the correct metal–ligand bond lengths than the non-bonded models, closely resembling the reference QM values. This observation further confirmed that obtaining correct metal–ligand bond lengths with non-bonded models, i.e., by adjusting only Lennard-Jones parameters, is challenging, particularly for coordination sites that include different ligands, as ZF1 and ZF2 of Npl4. However, with the bonded parameters for Cu^2+^, we still observed a slight preference to assume tetrahedral or vacant trigonal bipyramidal conformations for ZF1 and ZF2 over the seesaw conformation suggested by QM calculations. These discrepancies could be associated with the automatic parametrization method that we employed, based on the Seminario method, which is a fast and relatively simple approach and has approximations. To overcome these discrepancies, in future, other bonded models could be tested on Npl4 [35]. Furthermore, we could use more complex and accurate methods to obtain more accurate parametrization, including the definition of new atom types for all the relevant atoms in the metal site. Although these approaches do not allow for the study of dissociation events and changes in coordination geometry after metal replacement, they could provide insights into possible structural changes associated with the presence of Cu^2+^. Additionally, testing the performance of available polarizable force field and employing hybrid approaches based on QM/MM MD simulations, as was recently proposed for zinc metalloproteins [108], could be interesting avenues for our further investigation.

## Data availability

The scripts, input, and main outputs from this study will be made available in the GitHub repository associated with the publication https://github.com/ELELAB/NPL4_MD_QM_cupric_ions. The trajectories, QM calculation outputs and other large files are deposited in the OSF repository https://osf.io/4×5ye/

## Highlights

- The metabolite Cu^2+^ ions from anti-cancer Antabuse target NPL4 zinc fingers
- Assessment of MD force-field parameters for Zn^2+^ /Cu^2+^ on constructs of yeast Npl4
- Comparison with quantum mechanics of the metal coordination site of Npl4
- Non-bonded parameters fail to preserve metal coordination geometries
- Developed bonded parameters to treat Cu^2+^ and metal-coordinating atoms in Npl4

