## Supplementary figures and images for "Comparison of force fields to study the zinc-finger containing protein NPL4, a target for Antabuse in cancer therapy"

### Figure S1

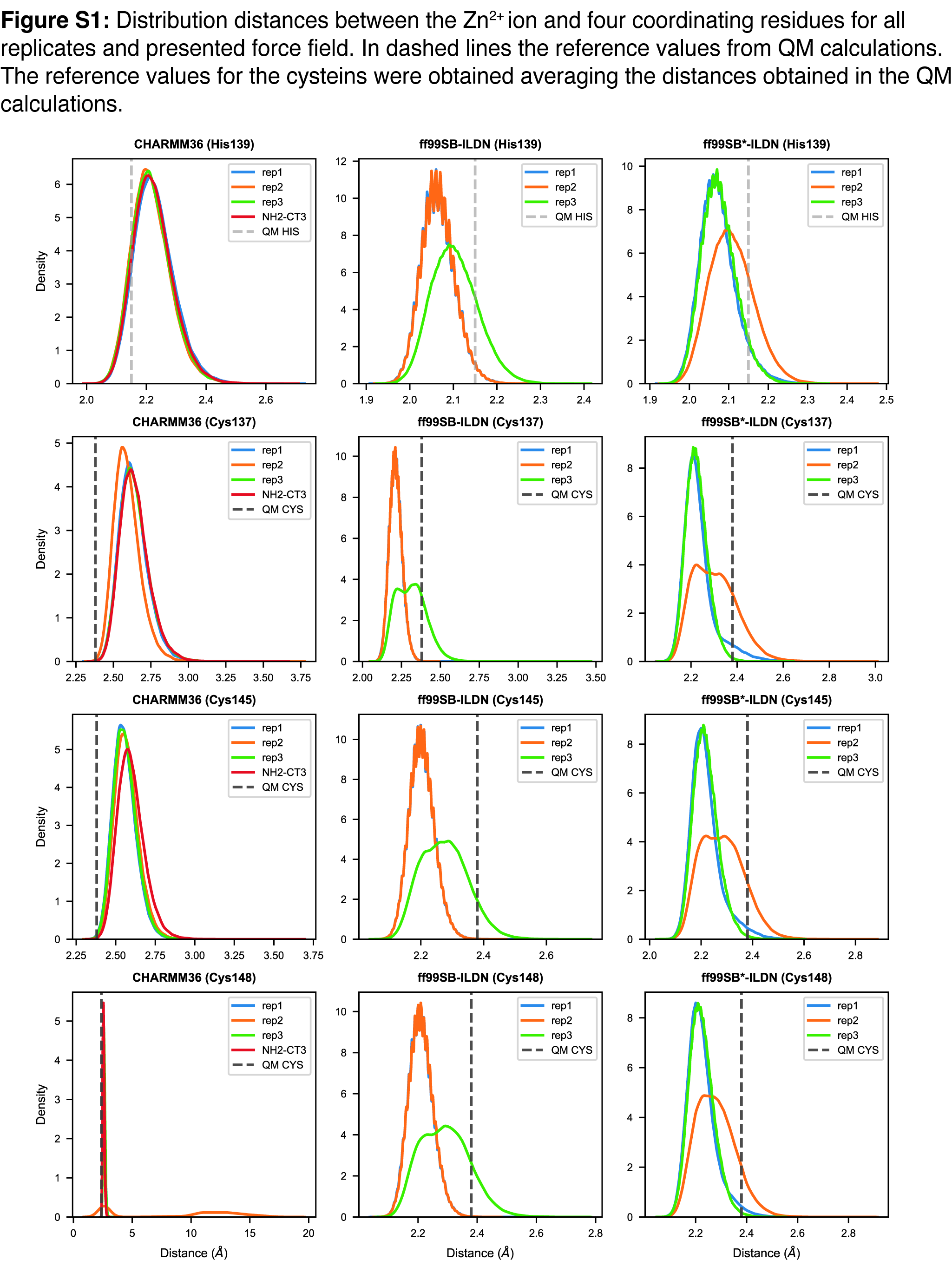

### Figure S2

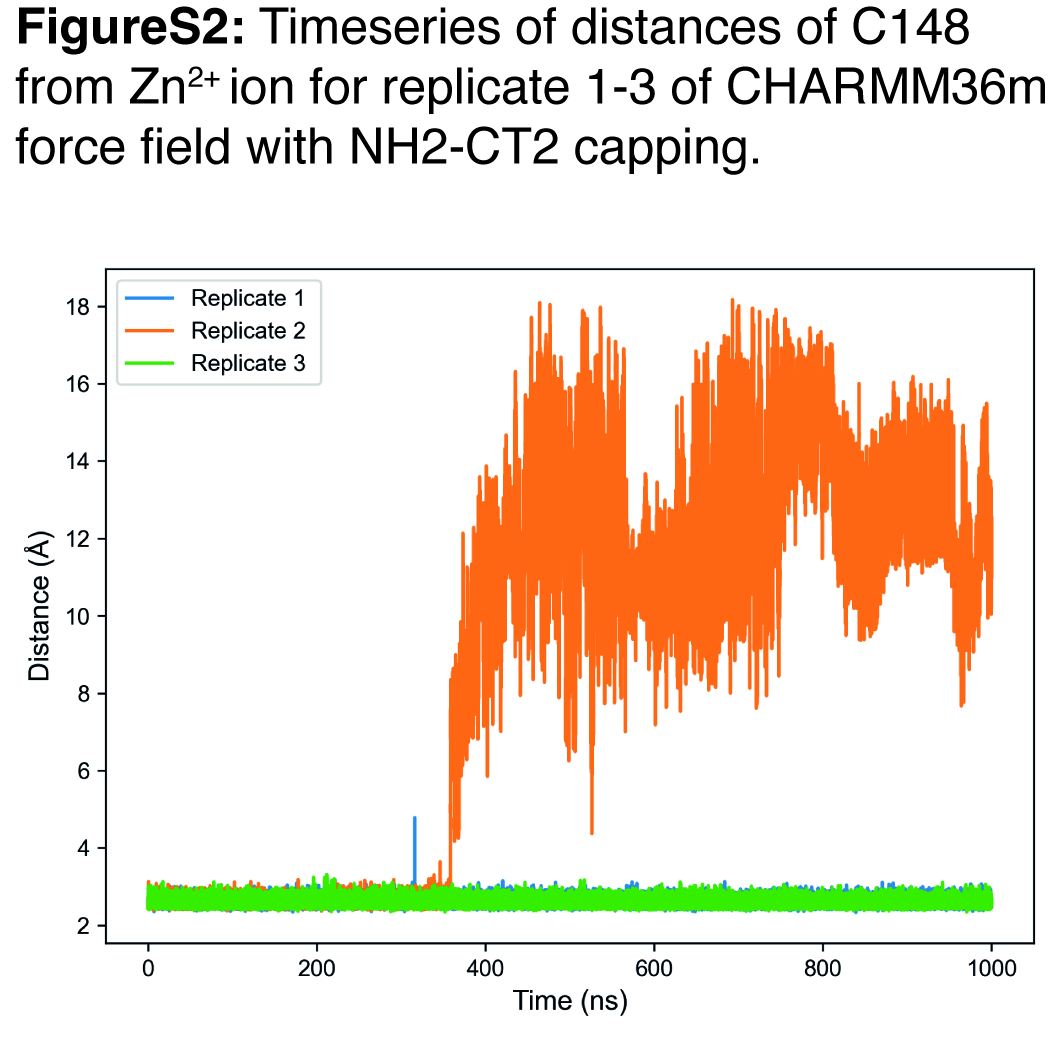

### Figure S3

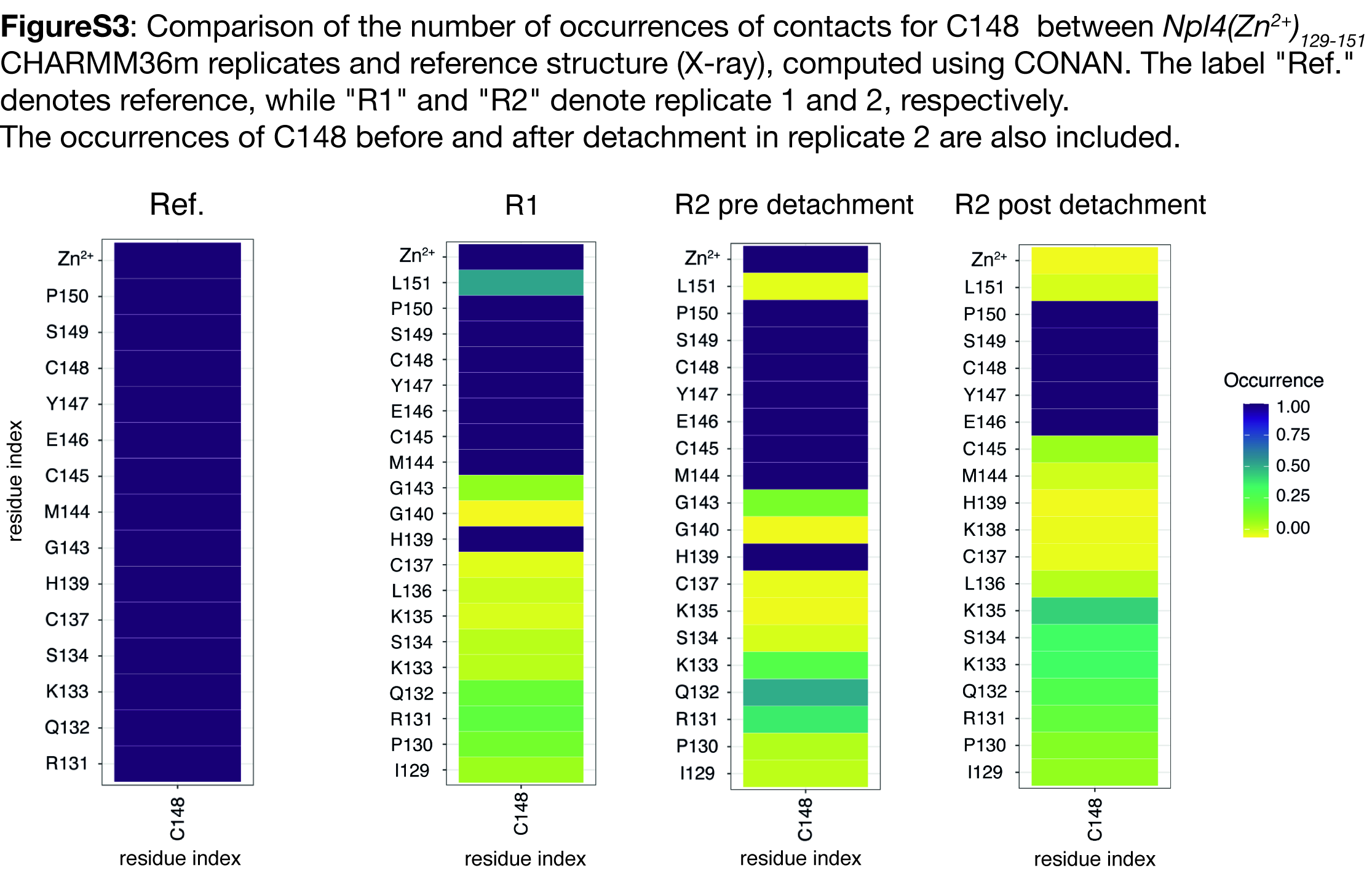

### Figure S4

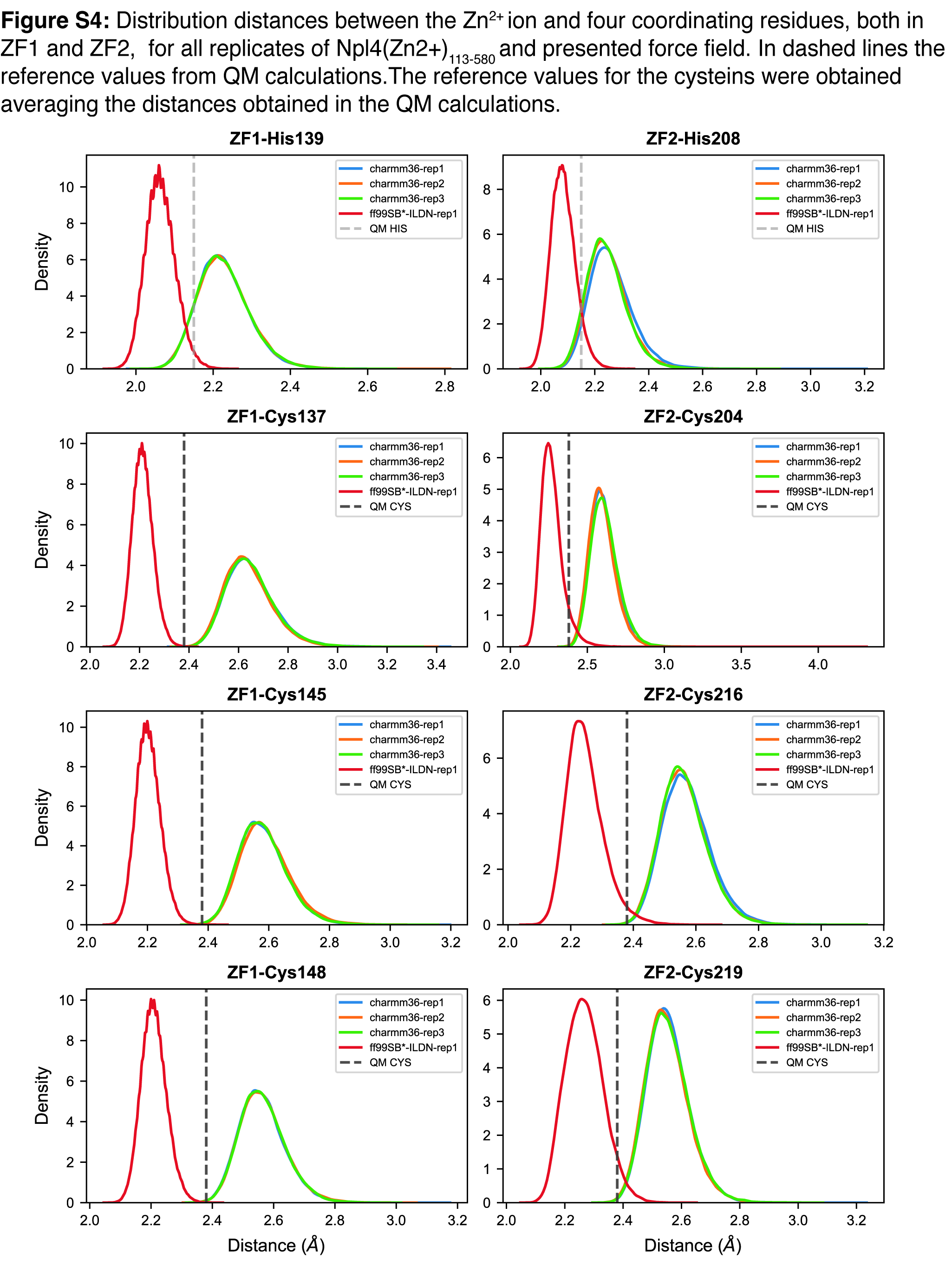

### Figure S5

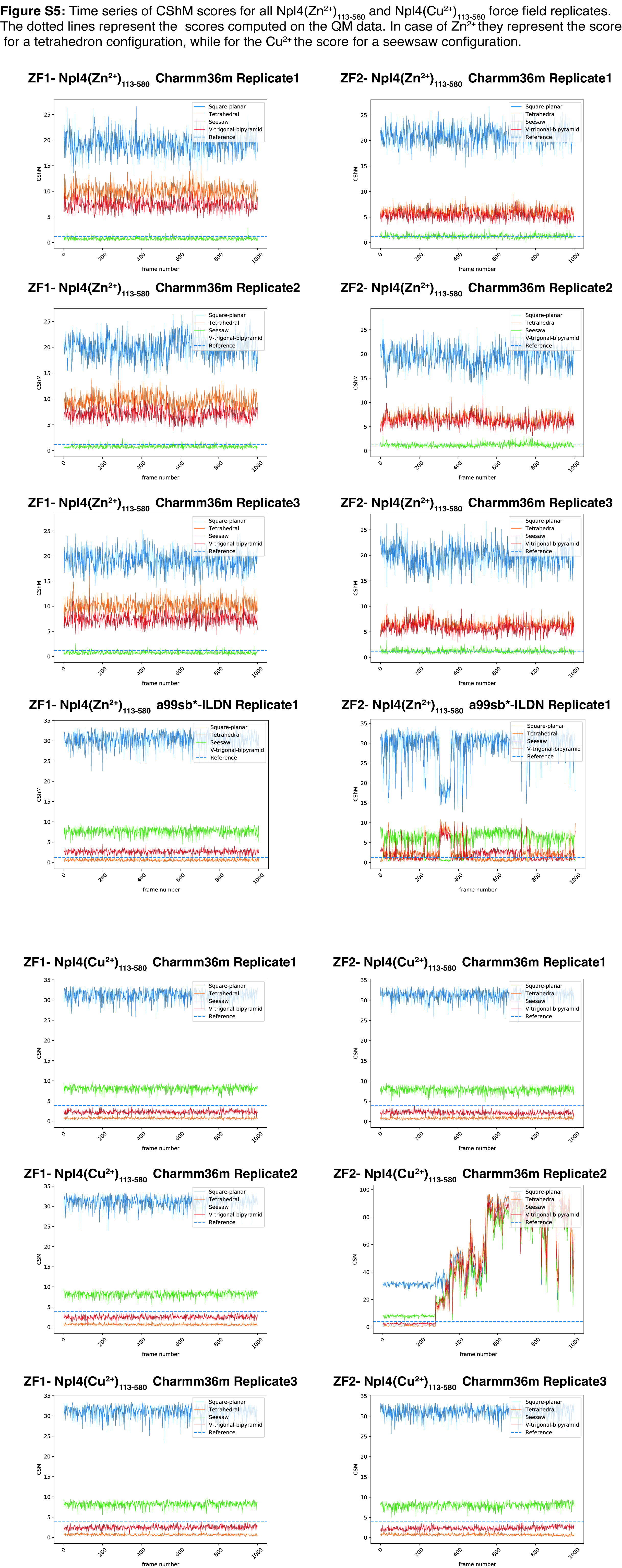
