## Supplementary material for "Comparison of force fields to study the zinc-finger containing protein NPL4, a target for Antabuse in cancer therapy": Table S1

| **System Details** | | | | | | | | | **Average CShM score** | | | |
| --- | --- | --- | --- | --- | --- | --- | --- | --- | --- | --- | --- | --- |
| **Name** | | **Force Field** | | **Capping** | | **Temperature (K°)** | | **n° replicate** | **Square (SP-4)** | **Tetrahedron (T-4)** | **Seesaw (SS-4)** | **Vacant trigonal bipyramid (vTBPY-4)** |
| Npl4(Zn^2+^)_129-151_ | | Charmm36 | | NH2-CT2 | | 298 | | 1 | 19.36 | 9.73 | **0.72** | 7.21 |
| Npl4(Zn^2+^)_129-151_ | | Charmm36 | | NH2-CT2 | | 298 | | 2 | 35.65 | 34.38 | 25.47 | 30.74 |
| Npl4(Zn^2+^)_129-151_ | | Charmm36 | | NH2-CT2 | | 298 | | 3 | 19.22 | 10.05 | **0.77** | 7.34 |
| Npl4(Zn^2+^)_129-151_ | | Charmm36 | | NH2-CT3 | | 298 | | 1 | 20,08 | 9.54 | **0.77** | 7.00 |
| Npl4(Zn^2+^)_129-151_ | | ff99sb-ILDN | | ACE-NH2 | | 298 | | 1 | 30.29 | **0.57** | 7.65 | 2.59 |
| Npl4(Zn^2+^)_129-151_ | | ff99sb-ILDN | | ACE-NH2 | | 298 | | 2 | 30.08 | **0.58** | 7.61 | 2.64 |
| Npl4(Zn^2+^)_129-151_ | | ff99sb-ILDN | | ACE-NH2 | | 298 | | 3 | 22.69 | 5.20 | **3.08** | 5.05 |
| Npl4(Zn^2+^)_129-151_ | | ff99sb*-ILDN | | ACE-NH2 | | 298 | | 1 | 28.76 | **1.51** | 6.68 | 3.05 |
| Npl4(Zn^2+^)_129-151_ | | ff99sb*-ILDN | | ACE-NH2 | | 298 | | 2 | 23.94 | 4.89 | **3.42** | 4.57 |
| Npl4(Zn^2+^)_129-151_ | | ff99sb*-ILDN | | ACE-NH2 | | 298 | | 3 | 30.39 | **1.24** | 7.20 | 2.00 |
| **System Details** | | | | | | | | | **Average CShM score (ZF1 / ZF2 )** | | | |
| **Name** | **Force Field** | | **Capping** | | **Temperature (K°)** | | **n° replicate** | | **Square (SP-4)** | **Tetrahedron (T-4)** | **Seesaw (SS-4)** | **Vacant trigonal bipyramid (vTBPY-4)** |
| Npl4(Zn^2+^)_113-580_ | | Charmm36m | | NH2-CT2 | | 298 | | 1 | 19.24 / 20.92 | 10.07 / 6.11 | **0.74 / 1.23** | 7.32 / 5.38 |
| Npl4(Zn^2+^)_113-580_ | | Charmm36m | | NH2-CT2 | | 298 | | 2 | 19.95 / 19.53 | 9.59/ 6.68 | **0.77 / 1.20** | 6.97 / 6.10 |
| Npl4(Zn^2+^)_113-580_ | | Charmm36m | | NH2-CT2 | | 298 | | 3 | 19.12 / 19.64 | 10.13 / 6.47 | **0.73 /1.19** | 7.40 / 5.84 |
| Npl4(Zn^2+^)_113-580_ | | ff99sb*-ILDN | | ACE-NH2 | | 298 | | 1 | 30.29 / 29.015 | **0.57** / 2.44 | 7.65 / 5.80 | 2.60 / **2.17** |
| Npl4(Cu^2+^)_113-580_ | | Charmm36m | | NH2-CT2 | | 298 | | 1 | 31.17 / 31.17 | **0.74 / 0.83** | 8.09 / 7.81 | 2.25 / 2.17 |
| Npl4(Cu^2+^)_113-580_ | | Charmm36m | | NH2-CT2 | | 298 | | 2 | 31.07 / 57.64 | **0.64 / 50.43** | 8.17 / 45.26 | 2.48 / 46.95 |
| Npl4(Cu^2+^)_113-580_ | | Charmm36m | | NH2-CT2 | | 298 | | 3 | 31.01 / 31.029 | **0.65 / 0.74** | 8.16 / 7.98 | 2.46 / 2.36 |
| Npl4(Cu^2+^)_113-580_ | | ff99sb*-ILDN bonded | | ACE-NH2 | | 298 | | 1 | 25.94 / 28.02 | **1.86 / 2.06** | 5.49 / 5.35 | 3.36 / 2.48 |

| **System Details** | | | | | **Average CShM score (ZF1 / ZF2)** | | | | |
| --- | --- | --- | --- | --- | --- | --- | --- | --- | --- |
| **Name** | **Force Field** | **Capping** | **Temperature (K°)** | **n° replicate** | **Hexagon** | **Pentagonal pyramid** | **Octahedron** | **Trigonal prism)** | **Johnson pentagonal pyramid** |
| Npl4(Zn^2+^)_129-151_ | Charmm36 | NH2-CT2 | 298 | 1 | 32.17 | 26.86 | 1.55 | 15.08 | 29.94 |
| Npl4(Zn^2+^)_129-151_ | Charmm36 | NH2-CT2 | 298 | 2 | 45.38 | 38.64 | 31.78 | 34.35 | 39.40 |
| Npl4(Zn^2+^)_129-151_ | Charmm36 | NH2-CT2 | 298 | 3 | 32.23 | 26.79 | 1.56 | 15.04 | 29.81 |
| Npl4(Zn^2+^)_129-151_ | Charmm36 | NH2-CT3 | 298 | 1 | 32.15 | 26.66 | 1.58 | 14.81 | 29.70 |
| Npl4(Zn^2+^)_113-580_ | Charmm36m | NH2-CT2 | 298 | 1 | 32.23 / 32.47 | 26.87 / 26.61 | 1.48 / 1.98 | 15.03 / 14.92 | 29.96 / 30.41 |
| Npl4(Zn^2+^)_113-580_ | Charmm36m | NH2-CT2 | 298 | 2 | 32.25 / 32.31 | 26.79 / 26.96 | 1.54 / 1.88 | 14.85 / 15.28 | 29.85 / 30.54 |
| Npl4(Zn^2+^)_113-580_ | Charmm36m | NH2-CT2 | 298 | 3 | 32.26 / 32.53 | 26.92 / 26.83 | 1.48 / 1.93 | 15.07 / 15.22 | 30.01 / 30.56 |
